# An open source MALDI-TOF mass spectrometry database for the identification of North American tick species of medical and veterinary importance

**DOI:** 10.1101/2025.06.19.660628

**Authors:** Ian Kirimi Daniel, Amber Holley, Samantha R. Hays, Pete D. Teel, Tanguy Tchifteyan, Guilherme G. Verocai, Maureen Laroche

## Abstract

Matrix-assisted laser desorption/ionization coupled with time-of-flight mass spectrometry (MALDI-TOF MS) is a fast emerging and robust method for rapid and accurate characterization of arthropod vectors. Correct identification of ticks is key in monitoring medically or veterinary relevant species and associated pathogens. It has been extensively used for tick identification based on leg proteins; however, in North America, MALDI-TOF MS has not been explored for comprehensive identification of ticks, except for a few studies focusing on a limited number of species. Herein, we have generated proteomic spectral profiles of nine North American tick species belonging to five genera obtained from laboratory-maintained colonies. From the 407 high-quality MS spectra generated in this study, 44 were deposited in our newly made database and used as reference. All remaining MS spectra were used to query and validate the database. All specimens were identified correctly to the species level using MALDI-TOF MS, with reliable Log Score Values (LSVs) ranging from 1.72 to 2.86, and median and mean values of 2.41 and 2.40, respectively. A single exception noted: an *Amblyomma mixtum* was misidentified as *Amblyomma tenellum*. As a cost-effective, user-friendly, and high-throughput method, MALDI-TOF MS can serve as an alternative for accurately identifying tick species of veterinary and public health importance in North America, thereby supporting tick-borne disease surveillance efforts.

## 1. Introduction

Ticks are important ectoparasites and vectors of veterinary and medically significant pathogens. While mosquitoes are the number one vectors in transmitting vector-borne pathogens (VBPs) worldwide, ticks are the primary vectors in the United States of America (USA), accounting for approximately 77% of all VBPs (Petersen et al., 2018, Rosenberg, 2018). Resident species such as the lone star tick (*Amblyomma americanum)*, the Gulf Coast tick (*Amblyomma maculatum)*, and the blacklegged tick (*Ixodes scapularis)*, as well as the newly introduced Asian long-horned tick (*Haemaphysalis longicornis),* are expanding their geographic ranges across North America (Phillips and Sundaram, 2024, Molaei et al., 2022, McClung and Little, 2023). Rapid and accurate identification of such species is critical for effective tick and tick-borne disease surveillance and control (Phillips and Sundaram, 2024, Molaei et al., 2022, McClung and Little, 2023).

Taxonomic uncertainties caused by morphological and genetic similarities among tick taxa may lead to incorrect or partial species identification (Alfred and Mertins, 2022). Historically, tick identification has relied on morphological criteria, which requires intense entomological training and expertise (Dantas-Torres, 2018, Nava et al., 2014b) and may be limited by differences across developmental stages (Coley, 2015), species complexes (Latrofa et al., 2013), and specimen integrity due to damaged/distorted features associated with storage conditions, fixation methods, or physiological status of the tick. Molecular assays for the characterization of tick DNA sequences can be time-consuming, costly, and may be limited by lacking or inconclusive sequence data for many taxa in Genbank (Dantas-Torres, 2018). Morphological and molecular identification methods both face difficulties when challenged with closely related species (Durden and Beati, 2013), including taxa within species complexes (Nava et al., 2014b). Advanced identification of these species requires diagnostic tools with precision surpassing traditional morphological and molecular methods. Matrix-assisted laser desorption/ionization time-of-flight mass spectrometry (MALDI-TOF MS) has become a crucial diagnostic tool in medical and veterinary entomology (Laroche et al., 2017, Sevestre et al., 2021). The technique offers several key advantages: high throughput capabilities, cost-effective implementation for routine surveillance, a user-friendly operation, and standardized sample preparation protocols – that remain consistent across arthropod families (Yssouf et al., 2016, Laroche et al., 2017). Unlike molecular approaches that require species-specific or genus-specific primer selection, and optimization, MALDI-TOF MS employs universal protein extraction and analysis protocols, significantly streamlining identification workflows for diverse arthropod taxa (Sevestre et al., 2021).

MALDI-TOF MS has traditionally been used for the identification and analysis of large biomolecules, including proteins, peptides, and nucleic acids (Aebersold and Mann, 2003, Roepstorff, 2000). For entomological applications, samples are homogenized in a solvent composed of formic acid and acetonitrile, and co-crystallized with an acidic matrix. The proteins are then subjected to laser ionization, and their time-of-flight, which correlates to their mass-to-charge ratio, is measured by an MS detector. Protein detection is then reflected as peaks, which together compose a spectrum that is representative of the original sample. A spectral database is then built through a meticulous selection of high-quality spectra from formally identified samples (Yssouf et al., 2013a), which can be later used to identify uncharacterized samples. The development of this method for the identification of arthropod vectors has resulted in an increasing number of applications across various arthropod families in the last decade (Galletti et al., 2024, M’madi et al., 2024, Huynh et al., 2022, Ndiaye et al., 2022, Sánchez-Juanes et al., 2022, Sevestre et al., 2021).

North America hosts numerous medically and veterinary significant tick species that serve as vectors for a diverse range of pathogens (Beard et al., 2021, Sonenshine, 2018). The U.S. reports approximately 50,000 human tickborne disease cases annually to the Centers for Disease Control and Prevention (CDC), though actual cases are estimated to exceed 500,000 (Rosenberg, 2018). Lyme disease represents the predominant concern, accounting for over 80% of reported cases, followed by Rocky Mountain spotted fever, ehrlichiosis, anaplasmosis, and babesiosis (Eisen et al., 2017, Rosenberg, 2018). The expanding geographic ranges of several North American tick species, coupled with the emergence of novel pathogens and the introduction of invasive tick species such as *Haemaphysalis longicornis* (Rainey et al., 2018, Sonenshine, 2018), have intensified the need for rapid and accurate tick identification capabilities. Furthermore, the presence of cryptic species complexes among North American ticks, such as the *Amblyomma maculatum* group and the *Rhipicephalus sanguineus* complex, presents additional challenges for conventional identification methods (Jones et al., 2017, Lado et al., 2018). Therefore, our study aimed to address the significant knowledge gap in MALDI-TOF MS characterization of North American tick species, as previous research has predominantly focused on African and European specimens. We evaluated the efficacy of this technology for accurate identification and differentiation among nine tick species of significant medical and veterinary importance in North America.

## 2. Materials and methods

### 2.1. Sample acquisition and storage

Adult specimens of nine tick species, representing five genera common to North America, were obtained from various sources across the USA. Males and females of *A. americanum*, *A. maculatum* sensu stricto (s.s.), *A. mixtum*, *A. tenellum*, *D. variabilis*, *R. sanguineus* s.l., *I. scapularis*, and *Ixodes pacificus* were included, and only females of *H. longicornis* were available (Table 1). All specimens were obtained alive from laboratory colonies from the Centers for Disease Control (CDC) for distribution by BEI resources NIAID, NIH in Virginia (VA), the Texas A&M AgriLife Tick Research Laboratory in Texas (TX), and the Oklahoma State University Tick laboratory, Oklahoma (OK) between 2023 and 2024. Newly obtained samples were immediately immobilized at −80 °C upon arrival and kept frozen until use. Species identity was already pre- determined as these were laboratory-reared ticks. However, all specimens were morphologically assessed for confirmation of identification under a stereo microscope (Carl Zeiss Microscopy, LLC, NY, USA) at a magnification of 40x during dissection. Representative specimens were selected for photographs for record keeping purposes using a digital microscope (VHX-7100, Keyence, USA) (Supplementary file 1)

**Table 1:**
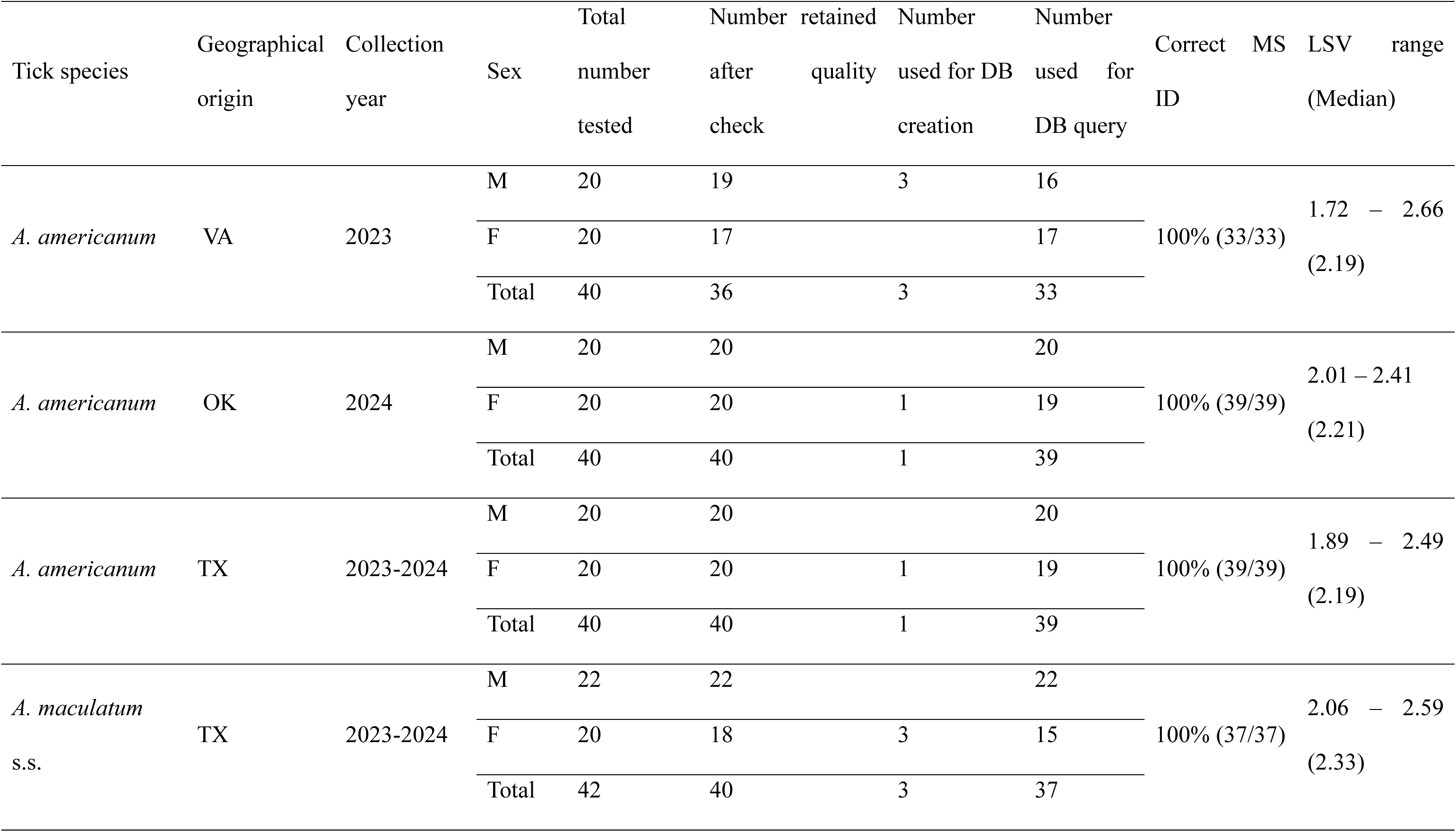
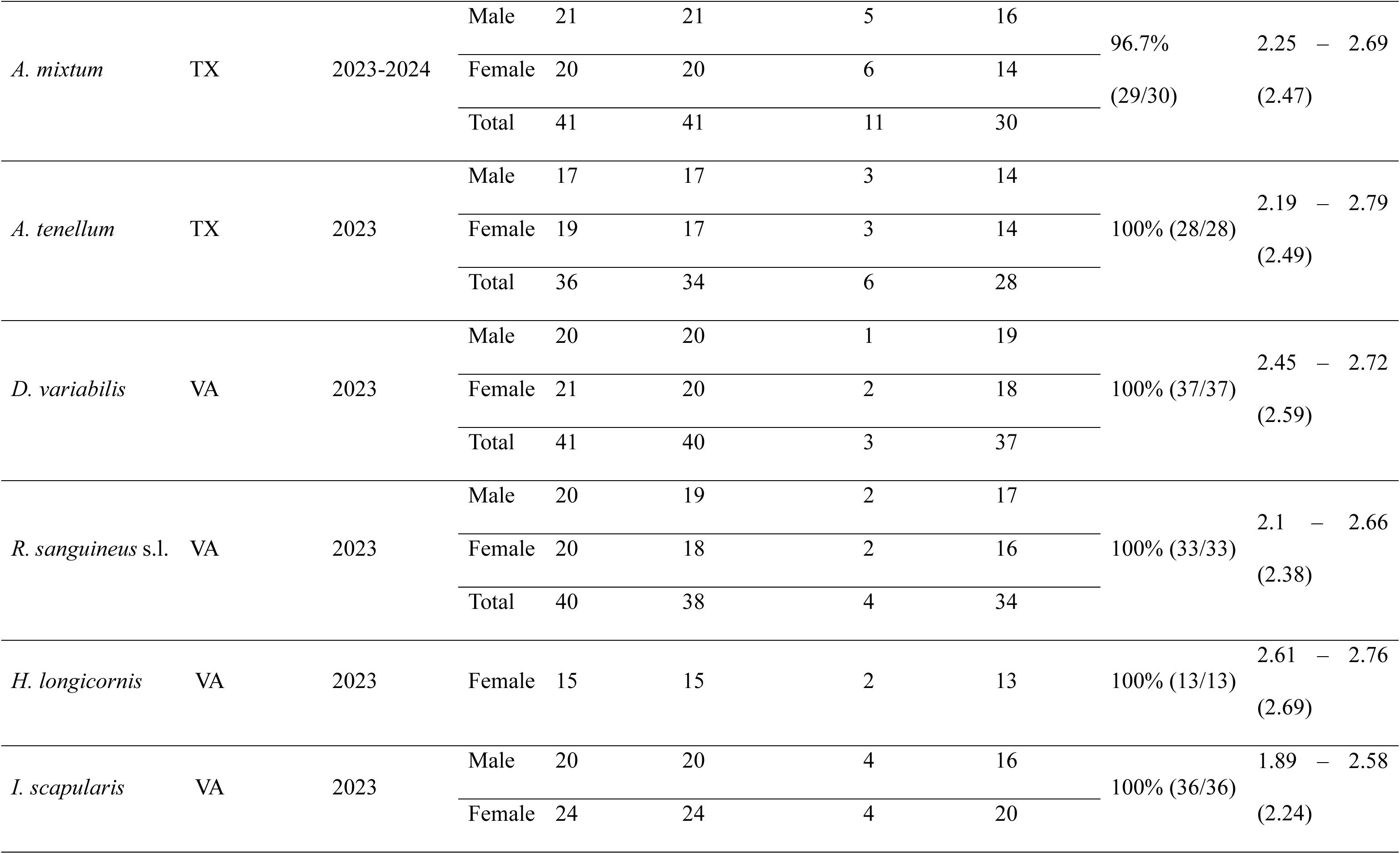
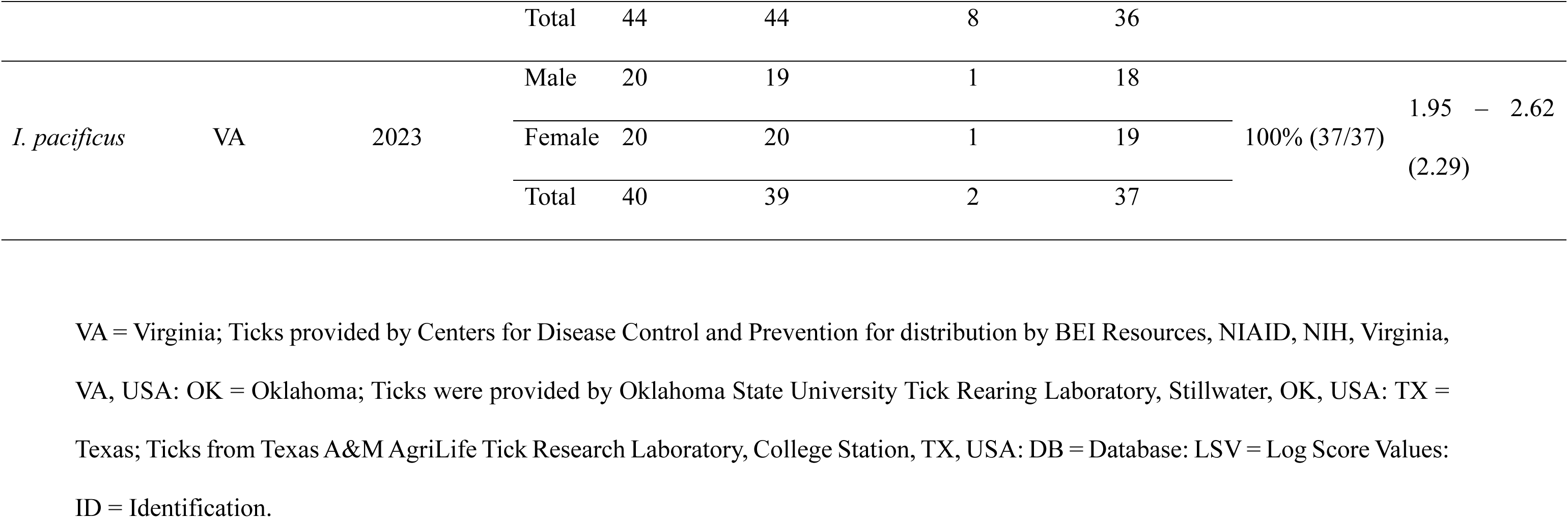
Results of MALDI-TOF MS identification for all North American tick species included in the study.

### 2.2. Sample preparation for MALDI-TOF MS analysis

Whole ticks were stored at −80 °C prior to use. Initially, specimens were briefly washed in 5% bleach for a few seconds, rinsed in distilled water, and carefully dried with a sterile paper towel. For each tick, four legs from one side were removed with sterile scalpel blades under a ZEISS Stemi 508 stereo microscope (Carl Zeiss Microscopy, LLC. NY, USA) at a magnification of 100x. The remaining body parts were stored at −20 °C as a backup sample while the removed legs were then pooled into an Eppendorf^®^ tube for MS analysis. To extract leg proteins, specimens were homogenized in a mix of 20 µL of 70% (v/v) formic acid (Sigma-Aldrich, Lyon, France) and 20 µL 50% (v/v) acetonitrile (Fluka, Buchs, Switzerland). Individual samples were then crushed in a

TissueLyser II device (Qiagen, Hilden, Germany) for three cycles of one minute per cycle at 30 Hz using glass powder (Glass beads, acid-washed G4649, ≤106μm, Sigma-Aldrich, Lyon. France) approximately equivalent to half the volume of the solution (i.e ∼20 µL displacement in a 40 µL solution). The homogenate was briefly centrifuged at 13000×g for one minute, and 1 μL of the final supernatant of each sample was spotted on an MSP 96 target polished steel target plate in quadruplicate (Bruker Daltonics, Wissembourg, France). After air drying, 1 μL of the matrix solution, composed of saturated α-cyano-4-hydroxycinnamic acid (Sigma-Aldrich, Lyon, France), 100% (v/v) acetonitrile, 100 % (v/v) trifluoroacetic acid (Sigma-Aldrich, Dorset, UK), and HPLC grade water, was overlaid on each sample. The matrix/analyte mix was co-crystallized at room temperature before being introduced to the MALDI-TOF MS instrument.

### 2.3. MALDI-TOF MS spectra analysis

We used a MALDI-TOF MS Microflex^®^ LRF mass spectrometer with the flexControl™ v.3.4 software (Bruker Daltonics, Bremen, Germany) to generate the protein spectra from each tick specimen. The flexControl™ software (Bruker Daltonics, Bremen, Germany) was set on a linear positive ion mode at a laser frequency of 60 Hz and the 2000 - 20000 m/z mass range. Each specimen-specific protein spectra corresponds to the ions accumulated from 240 laser shots taken randomly in 6 different positions of the same sample spot by automatic acquisition using the AutoXecute engine in Flex Control software (Bruker Daltonics, Bremen, Germany) at a maximum laser power of 40%. The resulting spectra were first visualized in FlexAnalysis version 3.0 (Bruker Daltonics, Bremen, Germany). The MS spectra were then exported to ClinProTools v2.2 and MALDI-Biotyper Compass Explorer v4.1.100 (Bruker Daltonics, Bremen, Germany) for data processing, including baseline subtraction, smoothing, and peak detection. For quality control purposes, the reproducibility and inter-species specificity of the MS spectra generated from the same tick and the same species were assessed by comparing the main spectrum profiles (MSP) using the MALDI-Biotyper Compass Explorer v4.1.100 software (Bruker Daltonics, Bremen, Germany). Specifically, the reproducibility and specificity of the spectra were evaluated using built-in statistical tests of the MS spectra, including principal component analysis (PCA) and cluster analysis (MSP dendrogram) with ClinProTools v2.2 and MALDI-Biotyper v4.1.100 software, respectively. ClinProTools and MALDI-Biotyper are widely used software from Bruker (Bruker Daltoniks, Bremen, Germany).

### 2.4. Processing of spectra and construction of the reference mass spectra database

We retained only the high-quality spectra of each species based on intra-species reproducibility, intensity (≥ 3000 a.u.), and absence of noise. These were used to create MSPs (Main Spectrum Profiles) using the MALDI-Biotyper Compass Explorer v4.1.100 software (Bruker Daltonics, Bremen, Germany), which are average spectra derived from all spots within a single sample. These MSPs indicate the most consistent spectral properties of a sample. Spectral selection for the database was supported using the MSP dendrogram function of the MALDI-Biotyper Compass Explorer (Bruker Daltonics, Bremen, Germany). MSP dendrograms are constructed based on a composite correlation index (CCI), a distance-based metric that clusters highly similar spectra. The CCI parameters quantify the spectral relationships by dividing spectra into intervals and comparing the correlation of these intervals across the dataset. The range of CCI values is from 0 to 1. A perfect correlation is denoted by a match value of 1, while 0 indicates the absence of correlation (Lawrence et al., 2019). Using dendrograms to select reference spectra ensures comprehensive coverage of the entire dataset. For a more comprehensive and unbiased query, we also incorporated previously generated reference spectra acquired by Dr. Maureen Laroche over the years in collaboration with other researchers. This in-house database contained ticks, lice, mosquitoes, and other arthropods from Africa and Europe and was used to assess potential crossmatch between the specimens in the database and the test tick spectra.

### 2.5. Database validation

The reliability of the database was validated through a query of the samples that were not used for database creation. Identification results were obtained with log score values (LSV), which indicate the confidence level of the identification. LSVs enable a thorough evaluation of reproducibility between a blind-tested spectrum and a reference spectrum, as they result from a detailed comparison of peak positions and intensities between these two spectra (Yssouf et al., 2013b). LSVs range from 0 to 3. A threshold of 1.8 was chosen for this study as it has been considered reliable for accurate arthropod species determination by MALDI-TOF MS in previous studies (Yssouf et al., 2013b). All LSVs were exported to Microsoft Excel for data cleaning and further processing.

## 3. Results

### 3.1. Spectral analysis

A total of 419 laboratory-reared ticks were subjected to MALDI-TOF MS analysis, representing nine species and five genera, namely, *Amblyomma* (4), *Dermacentor* (1), *Haemaphysalis* (1), *Ixodes* (2), and *Rhipicephalus* (1) (Table 1).

A total of 48 single mass spectra corresponding to 12 specimens were discarded after quality control, and the spectra obtained from 407 ticks were retained for database creation and query (Fig. 1).

**Figure 1:**
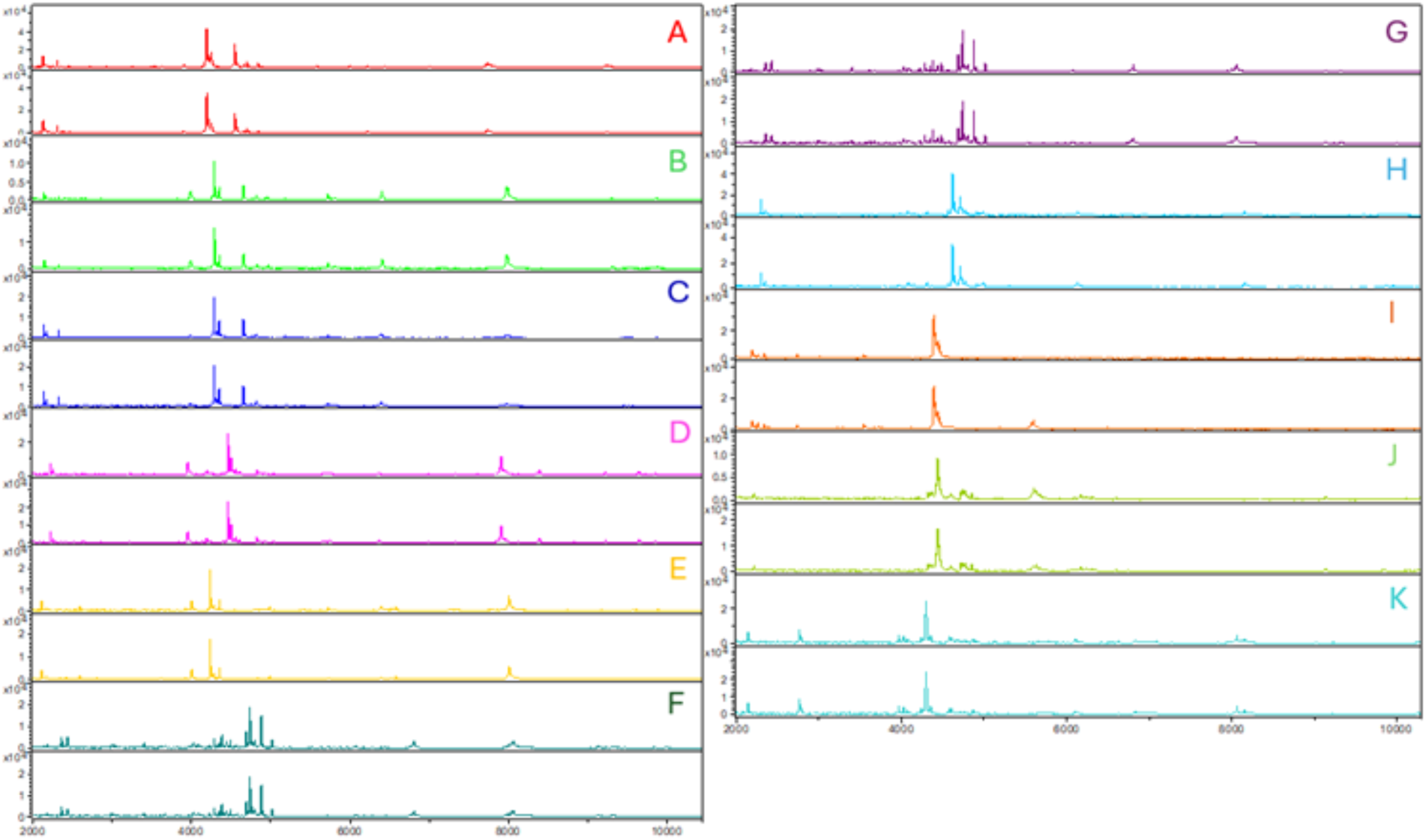
Representative MALDI-TOF mass spectral profiles of proteins from the legs of nine ixodid ticks. The intensity of spectra is measured in Arbitrary units (a.u.), and m/z indicates the mass-to-charge ratio as obtained using Flex analysis 3.0 software (Bruker Daltonics, Bremen, Germany). Each peak in the mass spectra signifies peptides or small proteins obtained from the tick legs. A) *Amblyomma americanum* (VA); B) *A. americanum* (OK); C) *A. americanum* (TX); D) *A. maculatum* s.s.; E) *A. mixtum*; F) *A. tenellum*; G) *Dermacentor variabilis*; H) *Haemaphysalis longicornis*; I) *Ixodes pacificus*; J) *I. scapularis*; K) *Rhipicephalus sanguineus* s.l.

A final dendrogram displaying hierarchical clustering using two to three representative specimens of each species confirmed the inter-species specificity across the dataset, and inter-population specificity across *A. americanum* samples from the three distinct laboratory colonies. Mixed clustering was observed for *A. mixtum* and *A. tenellum* (Fig. 2, Supplementary Material 1).

**Figure 2:**
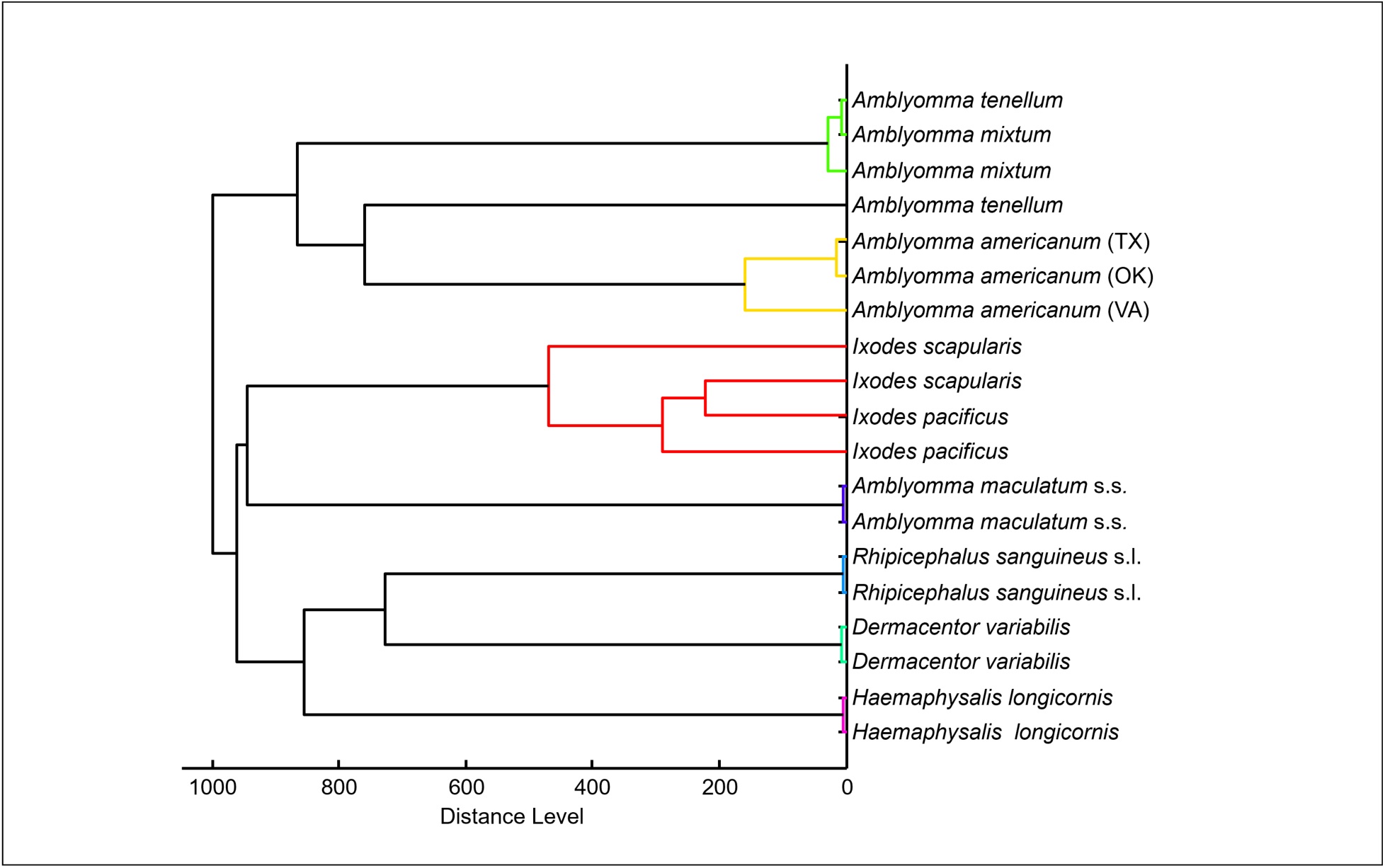
Dendrogram of the nine tick species. Cluster analysis of two to three representative spectral profiles per species, using MALDI Biotypher 3.0 software (Bruker Daltonics, Bremen, Germany). The distance is shown in relative units corresponding to relative similarity.

### 3.2. Database creation and validation

The database was built using representative spectra from two to eleven specimens from each tick colony, including *A. americanum* (5), *A. maculatum* s.s. (3), *A. mixtum* (11), *A. tenellum* (6), *D. variabilis* (3), *R. sanguineus* s. l. (4), *H. longicornis* (2), *I. scapularis* (8) and *I. pacificus* (2) (Table 1). Validation was performed by querying the remaining 363 specimens against this newly created database. The validation revealed that 96.8% to 100% of all specimens were correctly identified to the species level. The log score values ranged from 1.72 to 2.92 (mean = 2.44; median = 2.41; Table 1). Only one sample was identified with a score <1.8: a female *A. americanum* was correctly identified but with an LSV value of 1.72 (Table 1). *Amblyomma americanum* ticks from different colonies were identified correctly using reference spectra from a single colony; however, LSVs increased when a reference spectrum from each colony was added. A set of reference spectra developed in this study is available in the supplementary information of this manuscript for direct download (Supplementary file 2.0)

## 4. Discussion

Over the past decade, MALDI-TOF MS has remained a rapid, cost-effective, and relevant tool in microbiology and proteomics, and has been increasingly utilized in entomology and parasitology. Applications have expanded since 2019, and researchers continue to push the limits of integrative taxonomy to identify several species of arthropods such as bed bugs (M’madi et al., 2024), lice (Ouarti et al., 2020, Benyahia et al., 2020), mosquitoes (Huynh et al., 2022, Lawrence et al., 2019, Nebbak and Almeras, 2020), ticks (Galletti et al., 2024, Ahamada M’madi et al., 2022, Huynh et al., 2021), triatomine bugs (Laroche et al., 2018, Dos Santos Souza et al., 2019), and various dipterans (Silva et al., 2024, Arfuso et al., 2019). Once the device has been acquired, the technique offers cost-effective operation with minimal additional expenses per analysis (Laroche et al., 2017). While genomic sequencing of ticks may provide excellent sensitivity and specificity, the cost per sample is high ($10–$100) compared to ($0.43 −$3.59) for MALDI-TOF MS depending on the study (Yssouf et al., 2016, Laroche et al., 2017), which offers up to 87.8% savings in reagent and maintenance costs. Furthermore, molecular techniques are limited by the absence of universal primers for all tick species, lengthy nucleic acid extraction, and amplification protocols, and the lack of comprehensive reference sequences in the public repository (Yssouf et al., 2016, Diarra et al., 2017). Unlike morphological identification, which requires specialized expertise and is often limited by damaged specimens, immature stages, or cryptic species, MALDI-TOF MS provides rapid and accurate tick identification and maintains reliability even when specimens are physically compromised (Tran et al., 2015, Sung et al., 2018). Notably, some mass spectrometers such as the MALDI Biotyper Sirius (Bruker Scientific), a commonly available instrument in clinical microbiology laboratories, can process up to 96 samples in 10-15 minutes, making MALDI-TOF MS well-suited for high-throughput analyses (Seng et al., 2009).

North America hosts over 90 tick species of veterinary and public health importance, including vectors of Lyme disease, Rocky Mountain spotted fever, ehrlichiosis, and anaplasmosis (Eisen et al., 2017, Sonenshine, 2018). Despite this remarkable diversity and the medical significance of these arthropods, surprisingly few North American tick species have been characterized using MALDI-TOF MS. These include *Ixodes scapularis*, *A. americanum*, *Dermacentor variabilis* and *Dermacentor similis* (Galletti et al., 2024). Our spectral database achieved great identification accuracy (rates ranging from 96.7% to 100% across all species) and addresses a critical taxonomic gap by providing reference spectra for nine North American tick species: *I. scapularis*, *I. pacificus*, *A. americanum*, *A. maculatum* s.s., *A.mixtum*, *A. tenellum*, *D. variabilis*, *R. sanguineus* s.l., and the invasive *H. longicornis*.

Given the primary importance of Lyme disease as the most commonly reported vector-borne illness in the U.S., our database included the characterization of two *Ixodes* vectors. *Ixodes scapularis* and *I. pacificus* are vectors of critical medical importance due to their role in the transmission of *Borrelia burgdorferi* and *B. mayonii,* causative agents of Lyme disease, in North America (Boyer et al., 2019). In 2019, Boyer *et al*. demonstrated successful differentiation of closely related *Ixodes* species, including *I. scapularis* (with material from the University of Rhode Island, USA) and eight other *Ixodes* species from Europe (Boyer et al., 2019). However, Boyer *et al*. (2017) were unable to use MALDI-TOF MS to detect *B. burgdoferi* in *I. icinus*. Another study by Karger *et al*. (2012) characterized only the females of *I. scapularis* from a laboratory colony maintained in Berlin, Germany (Karger et al., 2012). The ability to distinguish between *I. scapularis* and *I. pacificus* is critical given their potentially varying vector competence for emerging pathogens like *Borrelia miyamotoi* and Powassan virus (Eisen et al., 2017). MALDI-TOF MS can differentiate immature stages of *Ixodes* ticks, which is crucial since nymphs are the primary vectors for human Lyme disease transmission and the most frequently encountered life stage during filed collections (Yssouf et al., 2013a, Karger et al., 2019). Future applications could extend to blood meal identification, as demonstrated in other arthropod vectors, with MALDI-TOF proving capable of identifying mixed blood meals and host sources in mosquitoes and sand flies (Hlavackova et al., 2019, Tuwei et al., 2025).

Beyond *Ixodes*, *Amblyomma* genus presents unique identification challenges and opportunities for MALDI-TOF MS diagnostics, particularly given the recent range expansion and emerging public health importance of several species in North America. *Amblyomma mixtum* and *A. tenellum* have faced a long-unresolved taxonomical discrimination. Indeed, *A. tenellum* has undergone repeated removal and reinstatement into the *A. cajennense* group over the years. Most recently, Beati et al. (2013) and Nava et al. (2014a) concluded that *A. tenellum* is a distinct species and not a synonym of *A. cajennense* (Nava et al., 2014b, Nava et al., 2014a, Beati et al., 2013). Genetic studies have also revealed that *Amblyomma imitator* (reclassified as a synonym of *A. tenellum* Koch, 1844) in the USA, is more closely related to *A. americanum* than to the *A. cajennense* group (Beati et al., 2013). MALDI-TOF MS has been shown to accurately discriminate mosquito and triatomine species complexes (Laroche et al., 2018, Huynh et al., 2022); however, challenges are always expected in developing highly specific spectra for cryptic species. The only misidentification encountered in our study was a male *A. mixtum,* which was incorrectly identified as *A. tenellum*. This challenge was anticipated due to the evolutionary relationships between these two species (Nava et al., 2014b, Beati et al., 2013).

The number of specimens used to build our database varied between two (*H. longicornis*) and eleven (*A. mixtum*) per species. This number varies based on the level of intra-species variability, which can result from population diversity (intrinsic or geographic), but also on the degree of inter-species specificity (i.e., a larger amount of data, herein spectra, is required to ensure differentiation of two closely related species) (Galletti et al., 2024, Boyer et al., 2019). In this study, a higher number of *A. mixtum* and *A. tenellum* was used to capture the similarities and specificities of these species.

Within the *Amblyomma* genus, *A. americanum* is a notable public health threat due to its aggressive host-seeking behavior and role as a vector for multiple pathogens, including *Ehrlichia chaffeensis* (human monocytic ehrlichiosis), *Francisella tularensis* (tularemia), and Heartland virus (Paddock and Childs, 2003, Beard et al., 2021, Goddard and Varela-Stokes, 2009). Additionally, its bites are associated with alpha-gal syndrome, an emerging IgE-mediated allergy to red meat (Commins et al., 2011). The species has undergone significant range expansion northward into northeastern states and westward into the Great Plains, establishing genetically distinct populations in previously non-endemic regions (Sonenshine and Roe, 2014, Springer et al., 2015, Fowler et al., 2022). Our MALDI-TOF MS analysis demonstrated distinct proteomic profiles for each colony, highlighting that MALDI-TOF MS can detect population-level differences. While this could be due to the fact that these colonies are originally maintained in different places, it could also reflect the known genetic diversity of geographically separated *A. americanum* populations (Martinez-Villegas et al., 2024, Monzón et al., 2016, Trout Fryxell et al., 2015). Indeed, MALDI-TOF MS was shown able to accurately distinguish populations of the same species of mosquitoes (Tandina et al., 2018, Fall et al., 2021), bed bugs (Benkacimi et al., 2020) and tsetse flies (Hoppenheit et al., 2014). These slight MS profile changes associated with geographic origin did not hinder species identification (Karger et al., 2019).

*Amblyomma maculatum* is another tick species with an expanding range in the United States. Once confined to the Gulf Coast states, *A. maculatum* has established populations as far north as Connecticut and as far inland as Illinois, with recent detection in urbarn epicenters including New York City and Philadelphia (Victoria et al., 2020, Molaei et al., 2024). This northward expansion coincides with remarkably high infection rates of *Rickettsia parkeri*, ranging from 20-56% in field-collected ticks (Fornadel et al., 2011, Varela-Stokes et al., 2011). Some populations in the northeast exhibit infection rates exceeding those in ticks from the historical range (Fornadel et al., 2011, Varela-Stokes et al., 2011). The ability of MALDI-TOF MS to accurately identify *A. maculatum* becomes critical as this species can also transmit *Hepatozoon americanum* and serve as a potential reservoir for other pathogens circulating in the environment (Ewing et al., 2002). Furthermore, *A. maculatum* presents unique diagnostic challenges as it belongs to a species complex that includes morphologically similar but genetically distinct species such as *A. triste, A. tigrinum*, and the recently described *A. maculatum* s.l. from West Texas to Sky Islands of Arizona (Lynn et al., 2024, Lado et al., 2018). Future work is needed to expand our database to include all cryptic species within the *A. maculatum* complex to enhance surveillance and vector control efforts.

The successful monitoring of invasive species represents another critical application of MALDI-TOF MS. The Asian long-horned tick, *H. longicornis*, is native to Asia but has spread to various continents, including North America. After the first encounter on a sheep in New Jersey in 2017 and in archival samples dating back to 2010 (Bajwa et al., 2024, Rainey et al., 2018), it has spread across the USA affecting livestock and wildlife in at least 21 states, including Washington, D.C., with the most recent sightings reported in Illinois (WKMS, 2024) and Oklahoma in 2024 (Myers and Scimeca, 2024). While *H. longicornis* can transmit various pathogens, including severe fever with thrombocytopenia syndrome virus (SFTSV) and *Rickettsia* species in Asia, its vector competence for North American pathogens is still being investigated, though it has been shown to be a competent vector for *Theileria orientalis* in U.S. cattle (Dinkel et al., 2021). The species poses a considerable threat to human and veterinary health due to its unique capabilities to reproduce parthenogenetically, its ability to invade new environments, and its use of numerous hosts, including humans, domesticated animals, and wildlife (Yabsley and Thompson, 2023). Morphological misidentification of *H. longicornis* is possible in areas where other *Haemaphysalis* species occur in North America, such as the rabbit tick *Haemaphysalis leporispalustris* (Thompson et al., 2020). Currently, MALDI-TOF MS spectra of *H. longicornis* are unavailable despite the urgent need to accurately detect and monitor this invasive non-native species and as new pathogens continue to be detected in this species (Molaei et al., 2025).

The incorporation of *Dermacentor variabilis* into our database builds upon recent advances in understanding the complex taxonomy of this genus (Galletti et al., 2024, Lado et al., 2021). In 2024, Galletti *et al*. developed the first nationwide MALDI-TOF mass spectrometry reference database for *Dermacentor* species in the U.S., making it publicly accessible through CDC MicrobeNet (https://microbenet.cdc.gov). This pioneering work applied MALDI-TOF MS to validate the distinction between closely related *D. variabilis and D. similis* using field-collected and archival museum specimens (Galletti et al., 2024). Databases for U.S. *Dermacentor* species are particularly valuable given the recent taxonomic reclassification by Lado *et al*. (2021), who demonstrated through integrative taxonomy that what was traditionally considered *D. variabilis* in the USA actually comprises two distinct species (*D. variabilis* and *D. similis*) delineated by geographic distribution and molecular markers (Lado et al., 2021). This widespread distribution underscores the importance of accurate identification, as these ticks represent the most widespread species parasitizing dogs and cats across the U.S (Duncan et al., 2021) and serves as key vectors for significant pathogens, including *Rickettsia rickettsii* and *Francisella tularensis*, and can cause tick paralysis in both humans and companion animals (Eisen et al., 2017). Our study complements Galletti et al.’s by utilizing laboratory-reared colonies to establish a database that includes protein profiles for both male and female *D. variabilis* specimens. Importantly, laboratory-reared specimens eliminate environmental contaminants and provide standardized references, while field-collected specimens like those used by Galletti *et al*. are a necessary validation to assess natural variation.

Field-caught samples are particularly relevant for species exhibiting high intra-specific variability, such as *Rhipicephalus* species (Boucheikhchoukh et al., 2018). Ticks belonging to the *R. sanguineus* group are important vectors of pathogens of medical and veterinary importance, including *Ehrlichia canis* (canine monocytic ehrlichiosis), *Babesia canis* and *B. vogeli* (canine babesiosis), *Hepatozoon canis*, and several rickettsiae (Dantas-Torres, 2008, Latrofa et al., 2014). In the United States, the role of brown dog ticks in the transmission of *R. rickettsi,* causative agent of Rocky Mountain spotted fever (RMSF), is of great concern, and the relative vector competence of *R. sanguineus* s.s. (aka, temperate lineage) and *R. linnaei* (aka, tropical lineage) is still unclear (Foley et al., 2025, Walker et al., 2022). The sympatric occurrence of *R. sanguineus* s.s and *R. linnaei* in Arizona where both lineages can infest the same dog simultaneously (Grant et al., 2023), presents significant diagnostic challenges as these lineages exhibit different vector competencies for *E. canis* (Villarreal et al., 2018). Several studies have confirmed that MALDI-TOF MS can identify *Rhipicephalus* ticks from various regions (Boucheikhchoukh et al., 2018). Using samples from European countries and Senegal, preliminary data from our group have shown that MALDI-TOF MS can distinguish *R. sanguineus* s.s. and *R. turanicus,* another species from the *R. sanguineus* group (Laroche et al., unpublished). Future studies must expand MALDI-TOF MS databases to include reference spectra from both *R. sanguineus* s.s and *R. linnaei*.

Finally making our reference spectra available in supplementary data, we aim to mitigate a current limitation of this approach. While more and more laboratories develop MS databases for medical and veterinary entomology, spectral libraries are rarely made publicly available. Although initiatives like MicrobeNet exist, they are still preliminary as they only include a limited amount of species and do not allow visualization of reference spectra. A publicly available and well-curated repository of spectral data for ixodid ticks would be of great value to the global scientific community.

## Conclusion

This work serves as a proof of concept for the use of MALDI-TOF MS in identifying North American ticks, and confirm its ability to identify cryptic tick species. However, we acknowledge the limitations of this study due to the lack of field-caught samples. Laboratory-reared specimens may not fully represent the protein profile variations found in wild populations but they serve as a robust foundation to establish standardized protocols that can be readily applied to field surveillance. Future studies should focus on validating this database with field-caught samples and evaluating the performance of MALDI-TOF MS to identify immature stages of species distributed in North America. Furthermore, the ability of MALDI-TOF MS to detect pathogens in these species should be evaluated.

## 6.2 Acknowledgments

The newly created reference spectra from this study were queried against an older database owned by Dr. Maureen Laroche. Several reference spectra from that database were created by Dr. Laroche when she was an employee of the Institut Méditerranée Infection (IMI) in Marseille, France. While Dr. Laroche is explicitly authorized to use these spectra for analysis, she does not own the intellectual property and cannot share the raw data. She thanks the IMI for letting her use these spectra for her current work.

The following reagents were provided by Centers for Disease Control and Prevention for distribution by BEI Resources, NIAID, NIH: Adult *Rhipicephalus sanguineus*, NR-42512; Live adult *Dermacentor variabilis*, NR-42513; Live adult *Amblyomma americanum*, NR-42514; Live adult *Ixodes pacificus* NR-44385 and live adult *Haemaphysalis longicornis*, NR-51846.

The following specimens were provided by Dr. Pete D. Teel at Texas A&M AgriLife Extension and the Department of Entomology at TAMU: Live adult *Amblyomma americanum*, Live adult *Amblyomma maculatum* sensu stricto, Live adult *Amblyomma mixtum*, and Live adult *Amblyomma tenellum*.

## 6.3. Data availability

Raw data corresponding to the reference spectra for each of the tick species used in this study are available in the attached supplementary file 2 to be downloaded freely. These files can only be opened and exported to a Bruker Scientific system.

## 6.3. Ethical approvals

There was no requirement for ethical approval in this laboratory-based study, as it did not involve any animal or human subjects.

## 6.3. CRediT authorship contribution statement

**Ian Kirimi Daniel:** Specimen acquisition, Writing – original draft, Writing – review & editing, Methodology, Formal analysis, Data curation, Visualization, Conceptualization. **Amber Holley:** Methodology, Data Curation, Formal Analysis, Visualization, Writing – review & editing. **Samantha Hays:** Sample acquisition, Writing – review. **Pete D. Teel:** Sample acquisition, Writing – review-. **Tanguy Tchifteyan**: Sample identification, imaging. **Guilherme G. Verocai:** Sample acquisition, Writing – review & editing, Conceptualization, Supervision. **Maureen Laroche:** Funding, Sample acquisition, Methodology, Data Curation, Formal Analysis, Data Validation, Conceptualization, Supervision, Writing – review & editing.

