## Supplementary figures and images for "An open source MALDI-TOF mass spectrometry database for the identification of North American tick species of medical and veterinary importance"

### Supplementary data 1

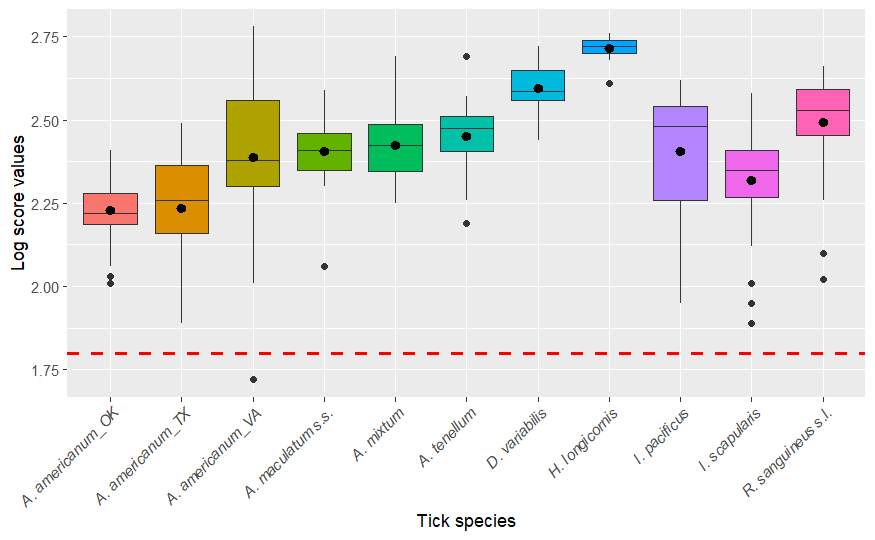
